# Evidence of polygenic adaptation at height-associated loci in mainland Europeans and Sardinians

**DOI:** 10.1101/776377

**Authors:** Minhui Chen, Carlo Sidore, Masato Akiyama, Kazuyoshi Ishigaki, Yoichiro Kamatani, David Schlessinger, Francesco Cucca, Yukinori Okada, Charleston W. K. Chiang

## Abstract

Adult height was one of the earliest putative examples of polygenic adaptation in human. By constructing polygenic height scores using effect sizes and frequencies from hundreds of genomic loci robustly associated with height, it was reported that Northern Europeans were genetically taller than Southern Europeans beyond neutral expectation. However, this inference was recently challenged. Sohail et al. and Berg et al. showed that the polygenic signature disappeared if summary statistics from UK Biobank (UKB) were used in the analysis, suggesting that residual uncorrected stratification from large-scale consortium studies was responsible for the previously noted genetic difference. It thus remains an open question whether height loci exhibit signals of polygenic adaptation in any human population. In the present study, we re-examined this question, focusing on one of the shortest European populations, the Sardinians, as well as on the mainland European populations in general. We found that summary statistics from UKB significantly correlate with population structure in Europe. To further alleviate concerns of biased ascertainment of GWAS loci, we examined height-associated loci from the Biobank of Japan (BBJ). Applying frequency-based inference over these height-associated loci, we showed that the Sardinians remain significantly shorter than expected (~ 0.35 standard deviation shorter than CEU based on polygenic height scores, *P* = 1.95e-6). We also found the trajectory of polygenic height scores decreased over at least the last 10,000 years when compared to the British population (*P* = 0.0123), consistent with a signature of polygenic adaptation at height-associated loci. Although the same approach showed a much subtler signature in mainland European populations, we found a clear and robust adaptive signature in UK population using a haplotype-based statistic, tSDS, driven by the height-increasing alleles (*P* = 4.8e-4). In summary, by examining frequencies at height loci ascertained in a distant East Asian population, we further supported the evidence of polygenic adaptation at height-associated loci among the Sardinians. In mainland Europeans, we also found an adaptive signature, although becoming more pronounced only in haplotype-based analysis.

## Introduction

Because of the highly polygenic nature of many human complex traits, polygenic adaptation was thought to be one of the important mechanisms of phenotypic evolution in humans. Since each genetic locus contributes a small effect to complex traits, polygenic adaptation is expected to be different from the classic selective sweep, where a beneficial allele is driven to near-fixation in a population due to strong positive selection. In polygenic adaptation, only a subtle but coordinated allele frequency shift across loci underlying the selected trait is expected. In humans, height is one of the earliest putative examples of polygenic adaptation. (We note that adaptation, if present, could be due to a trait for which height is a proxy; it is the height-associated loci that exhibited signals of adaptation, and thus we are investigating if selection had been contributing to the height differences between populations, rather than whether height itself is under selection.) Northern Europeans are known to be taller than Southern Europeans on average (Bentham et al., 2016; Grasgruber, Cacek, Kalina, & Sebera, 2014). By evaluating the changes of allele frequencies at height-associated loci, either weighted or unweighted by the effect sizes on height, multiple studies have suggested polygenic adaptation in human height in European and other populations (Berg & Coop, 2014; Field et al., 2016; Guo et al., 2018; Robinson et al., 2015; Tucci et al., 2018; Turchin et al., 2012; Zoledziewska et al., 2015)

However, the adaptative signature at height-associated loci was recently called into question by two papers (Berg et al., 2019; Sohail et al., 2019). The authors of both papers found that the adaptive signature disappeared if GWAS summary statistics from UK Biobank (UKB) were used in the analysis. This suggested that previous studies of adaptation might have been misled due to the ascertainment of a set of height-associated loci with biased estimates of effect sizes; the aggregation of which across large number of height-associated loci led to the apparent difference in genetic height between Northern and Southern Europeans. It was suggested that the biased effect sizes were due to residual uncorrected stratification from large-scale consortium studies of human height such as the GIANT (Genetic Investigation of ANthropometric Traits) consortium (Wood et al., 2014), where the control for population stratification occurred at the level of smaller individual studies were insufficient. In contrast, large-scale biobank level studies where individual data were available permitted much more effective control for stratification either through principal component analysis or linear mixed models (Berg et al., 2019; Sohail et al., 2019).

While these studies and others (Kerminen et al., 2019) investigated the degree to which over-estimated effect sizes in GWAS led to unrealistic polygenic height scores and differences between populations, it remains an open question of whether height-associated loci exhibit signals of polygenic adaptation, in any human populations. For one, the original report of polygenic adaptation on height in Europe relied solely on frequency changes between populations and the direction of association among alleles most associated with height (Turchin et al., 2012). By not taking into account the effect sizes this approach should be more robust to uncorrected stratification in GWAS. Moreover, loci most strongly associated with height appear to still exhibit a strong signal in haplotype-based singleton density score (SDS) analysis (Sohail et al. 2018). Furthermore, estimated temporal trajectory of polygenic height scores also showed a small but significant uptick in the recent history (Edge and Coop 2019). Finally, pygmies from the Indonesian island Flores exhibited lower genetic height than expected, based on height loci ascertained in the distantly related UKB population (Tucci et al., 2018).

In the present study, we re-examined if height-associated loci exhibit signs of adaptation in Europe. We characterized the behavior of polygenic adaptation analysis and how the conclusions and interpretations are impacted by different ascertainment schemes in the analysis. Moreover, in light of reported stratification (Haworth et al., 2019) and polygenic selection for height (Liu et al., 2018) in the UKB population, and our finding here that height-associated loci ascertained from UKB are still significantly associated with structure in Europe, we chose to conduct our analysis using height-associated SNPs ascertained from the Biobank of Japan (BBJ). Because it is a population distant from Europe, we reasoned and demonstrated that height-associated loci ascertained in this panel were not associated with the structure in Europe. As such, in absence of a very large scaled family-based analysis to ascertain height-associated loci, our analytical framework represents the least likely to be impacted by any cryptic covariances due to population stratification. Under this framework, we found that the Sardinians, one of the shortest populations in Europe, have significantly lower polygenic height scores than expected given their genetic relatedness to other European populations, consistent with previous reports (Zoledziewska et al., 2015). In mainland Europe, however, the adaptive signature based on allele frequencies was much subtler, though we observed a strong haplotype-based signature using tSDS (trait-SDS). Together, findings of our study provide additional evidence of polygenic adaptation at height-associated loci in at least some European populations.

## Materials and methods

### GWAS panels

To calculate polygenic height score, we obtained GWAS summary statistics from three studies.

- GIANT (Wood et al., 2014): a meta-analysis of 79 separate GWAS for height involved a total of ~253 K individuals of European ancestry with ~2.5 M variants, with each study independently controlling for population structure via the inclusion of principal components as covariates.
- UKB: a GWAS on UKB with ~360 K individuals, imputed to whole-genome sequencing data from Haplotype Reference Consortium (HRC), UK10K, and 1000 Genomes for ~13.8 M variants, corrected for 20 principal components (https://storage.googleapis.com/ukbb-robert/height_ukb_giant/robert1/50.imputed_v3.results.both_sexes.tsv.gz downloaded in October 2018).
- BBJ (Akiyama et al., in press): ~159 K individuals of Japanese ancestry from Biobank of Japan (Hirata et al., 2017; Nagai et al., 2017), imputed to combined whole-genome sequencing data from BBJ and 1000 Genomes for 27.2 M variants. Individuals not of Japanese origins were excluded by self-report or principal component analysis (PCA). GWAS was conducted using linear mixed model implemented in BOLT-LMM (P.-R. Loh et al., 2015).

### Population genetic data

We separately evaluated polygenic selection on height-associated loci in mainland Europeans and in Sardinians. For mainland Europeans, we analyzed two populations from Northern Europe, *i.e*. GBR (British in England and Scotland) and CEU (Utah Residents with Northern and Western European Ancestry), and two populations from Southern Europe, *i.e*. IBS (Iberian Population in Spain) and TSI (Toscani in Italia) using data from 1000 Genomes phase three release (1000 Genomes Project Consortium et al., 2015). We did not include the FIN (Finnish in Finland) population because of its known unique demographic history (Locke et al., 2019), and for achieving better balance of sample sizes between the two comparison populations. For Sardinians, we further included frequency estimates from 615 Sardinian individuals whole-genome sequenced in the SardiNIA study (Chiang et al., 2018) along with the four mainland European populations.

### Population structure analysis

To measure the impact of uncorrected stratification on estimated effect sizes for a set of ascertained height-associated variants, we computed the correlation between principal component (PC) loadings and beta effects estimated from GWAS. We first conducted PCA on the four mainland European populations (CEU, GBR, IBS, and TSI) from 1000 Genomes. We used variants that were present in all three GWAS panels, and that had minor allele frequency (MAF) > 5% in the four European populations. We pruned SNPs with r^2^ < 0.2 with the option of ‘--indep-pairwise 50 5 0.2’ using PLINK version 1.9 (www.cog-genomics.org/plink/1.9/) (Chang et al., 2015) to remove correlated variants, and removed SNPs in regions of long-range LD (Price et al., 2008). PCA was performed on the remaining variants using Eigensoft version 7.2.1 (https://github.com/DReichLab/EIG/archive/v7.2.1.tar.gz). We performed linear regressions of the PC scores on the allelic genotype count for each variant and used the resulting regression coefficients as the variant’s PC loading estimates. We then computed Pearson correlation coefficients of PC loadings and beta effects from each GWAS panel (GIANT, UKB, and BBJ) for each principal component, and estimated *P* values based on Jackknife standard errors by splitting the genome into 1,000 blocks with an equal number of variants.

### Population-level polygenic height score calculation

To compute polygenic scores, we ascertained independent GWAS variants associated with height by selecting a set of genome-wide significant variants (p < 5e-8) with MAF > 1% after greedily pruning any other variants such that no two variants were within 500 kb of each other. For simplicity, we refer to these sets of variants as independent, although they are only quasi-independent as linkage disequilibrium (LD) information was not taken into account. In order to compare to previous reports (Berg et al., 2019; Sohail et al., 2019), we additionally ascertained height-associated SNPs by dividing the genome into approximately independent LD blocks computed in (Berisa & Pickrell, 2016) (for GIANT and UKB panels, using the ~1,700 blocks in the EUR population; for BBJ panel, using the ~1,400 blocks in the ASN populations), and the variant with the lowest P value, regardless of whether the variant reached genome-wide significance level, within each block was ascertained to give a set of ~1,700 or 1,400 independent variants for each height GWAS panel.

Given a set of height-associated SNPs, the estimated effect sizes from each GWAS panel were then used to compute polygenic height scores for each population by

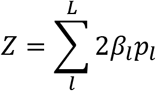

Where *p_l_* and *β_l_* were the allele frequency and effect size at SNP *l*. We would also compute the per-SNP contribution to differences between populations 1 and 2 as (*Z*_2_ – *Z*_1_)/*L*.

### Evaluation of selection on height-associated loci

To evaluate the evidence of selection on height-associated loci, we applied the following three methods:

#### Excess variance test

We conducted the Q_x_ test (Berg & Coop, 2014) to determine whether the estimated polygenic scores exhibited excess variance among populations than null expectation under genetic drift:

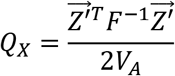

Where 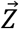 was a vector of estimated genetic values (*i.e*. a sum of sample allele frequencies weighted by effect size) for testing populations, *F* was a matrix describing the correlation structure of allele frequencies across populations, *V_A_* was the additive genetic variance of the ancestral (global) population. The *F* matrix was constructed by sampling 20,000 variants from GWAS panels matching by MAF, recombination rate, and background selection measured by B values from (McVicker, Gordon, Davis, & Green, 2009). Specifically, we partitioned variants into a three-way contingency table in each GWAS panel, with 25 bins for MAF (i.e. a bin size of 0.02, matched by ancestry), 100 bins for recombination rate, and 10 bins for B value. For recombination rate, we used CEU, GBR, and JPT (Japanese in Tokyo, Japan) genetic maps generated from the 1000 Genome phased OMNI data (http://ftp.1000genomes.ebi.ac.uk/vol1/ftp/technical/working/20130507_omni_recombination_rates/ downloaded in November 2018) for GWAS panels GIANT, UKB, and BBJ, respectively. The statistic should follow a *χ*^2^ distribution with *M-1* degrees of freedom under neutrality, where *M* was the number of test populations, from which an asymptotic p value was estimated. Significant excess of variance among populations would be consistent with the differential action of natural selection among populations.

To identify outlier populations which contributed to the excess of variance, we further estimated the conditional Z-score proposed by (Berg & Coop, 2014). Specifically, we excluded one population at a time, and then calculated the expected mean and variance of genetic value in the excluded population given the values observed in the remaining populations, and the covariance matrix relating them. Using this conditional mean and variance, we calculated a Z-score to describe the fit of the estimated genetic value of the excluded population by the drift model conditioned on the values in the remaining populations. An extreme Z-score for a particular population suggested that the excluded population had experienced directional selection on the trait of interest that was not experienced by the related populations on which we conditioned the analysis.

In practice, we also generated the empirical null distributions of Q_x_ statistic and conditional Z-scores by calculating 10,000 null genetic values using resampled SNPs genome-wide matching by MAF, recombination rate, and B value in the same way for the construction of *F* matrix. The empirical *p* values for conditional Z-scores tended to match well with the asymptotic *p* values (data not shown). Therefore, throughout the study, we used the asymptotic *p* value for the Q_x_ statistic and conditional Z-score. The scripts we used to implement these analyses are available at https://github.com/jjberg2/PolygenicAdaptationCode (downloaded in October 2018).

#### Polygenic height score trajectory

Based on the framework of (Edge & Coop, 2019), we constructed the history of polygenic height scores in the GBR and Sardinian populations. We extracted out 91 individuals each from GBR and Sardinia (Chiang et al., 2018), then used the software RELATE v1.0.8 (Speidel, Forest, Shi, & Myers, 2019) to reconstruct ancestral recombination graphs in these two populations together. We only included bi-allelic SNPs that are found in the genomic mask provided with the 1000 Genome Project dataset (downloaded from ftp://ftp.1000genomes.ebi.ac.uk/vol1/ftp/release/20130502/supporting/accessible_genome_masks/ in March 2019). We used an estimate of the human ancestral genome (downloaded from ftp://ftp.1000genomes.ebi.ac.uk/vol1/ftp/phase1/analysis_results/supporting/ancestral_alignments/ in March 2019) to identify the most likely ancestral allele for each SNP. We initially estimated branch lengths using a constant effective population size of 11,314 and a mutation rate of 1.25e-8 per base per generation. We then calculated mutation rate and coalescent rate through time given the branch lengths using default parameters (30 bins between 1,000 and 10,000,000 years before present, and 28 years per generation). By averaging coalescence rates over all pairs of haplotypes and taking the inverse, we obtained a population-wide estimate of effective population size. We then used this population size estimate to re-estimate branch lengths. We iterated these two steps five times to convergence, as suggested by (Speidel et al., 2019), then obtained a final estimate of branch lengths and the effective population size. Based on the output ancestral recombination graphs we estimated the time courses of polygenic height scores as estimated sum of allele frequencies weighted by effect sizes for GBR and Sardinia separately, using the three estimators proposed in (Edge & Coop, 2019): (1) the proportion-of-lineages estimator estimated allele frequency at a specified time in the past as the proportion of lineages that carry the allele of interest; (2) Waiting-time estimator and (3) lineages-remaining estimator estimated allele frequency as the relative sizes of the two subpopulations carrying ancestral and derived alleles. The former used waiting times between coalescent events to estimate subpopulation sizes, while the latter used the number of coalescence events that occur between specified time points. The same set of SNPs (genome-wide significantly associated variants in BBJ, after pruning by distance) was used to compute the polygenic height score in both GBR and Sardinia. We focused on the proportion-of-lineages estimators as it had been shown to be the most powerful at detecting selection, due to its improved precision (Edge & Coop, 2019), but all three estimators were provided for completeness.

We tested for significant differences in polygenic height score trajectory between GBR and Sardinia over time by performing 10,000 permutations of the signs of effect sizes across these SNPs. We specifically tested if polygenic height score in Sardinia is changing relative to GBR for two time intervals: between 20,000 years and 5,000 years before present, and between 5,000 years ago and the present time. These time points were chosen because they are approximately the time point for first evidence of human inhabitation on Sardinia (Calò, Melis, Vona, & Piras, 2008; Vona, 1997) and the onset of Neolithic period around which time Sardinia diverged from mainland Europeans (Chiang et al., 2018). To estimate courses of polygenic height scores and conduct significance test, we adopted code from (Edge & Coop, 2019) available at http://github.com/mdedge/rhps_coalescent (downloaded in April 2019).

#### tSDS analysis

We tested whether height-associated loci are under recent selection in a mainland European population by examining the distribution of tSDS. Recent selection results in shorter tip branches for the favored allele. The Singleton Density Score (SDS) (Field et al., 2016) leveraged the average distance between the nearest singletons on either side of a test SNP across all individuals to estimate the mean tip-branch length of the derived and ancestral alleles and used this measure to infer evidence of selection. The sign of SDS can be polarized such that positive scores indicate increased frequency of the trait-increasing (or trait-decreasing allele) instead of derived allele. The metrics is referred to as trait-SDS (tSDS). We obtained pre-computed SDS for 4,451,435 autosomal SNPs from 3,195 individuals from the UK10K project (downloaded from http://web.stanford.edu/group/pritchardlab/UK10K-SDS-values.zip in February 2019). In each GWAS panel (UKB and BBJ), we included only SNPs with a reported SDS prior to distance pruning to obtain a set of genome-wide significant SNPs. We then calculated the tSDS value for each variant by polarizing SDS to height-increasing alleles. Therefore, a positive tSDS indicates that a height-increasing allele has risen in frequency in the recent past; a negative tSDS indicates a height-increasing allele has dropped in frequency in the recent past. To estimate if an observed mean tSDS across a set of height-associated SNPs was significantly different from the null expectation, we performed 100,000 permutations of the sign of tSDS across these SNPs and reported the empirical *p*-value.

## Results

### European population structure underlying GWAS summary statistics

Incomplete control of population structure could lead to biases in the estimate of the effect sizes in GWAS, and as a result, polygenic scores constructed based on associated loci ascertained from these GWAS would show elevated population differentiation relative to neutral genetic drift (Robinson et al., 2015). For example, because the primary feature of genetic differentiation in mainland Europe is along the north-south axis, if human height is differentiated along this axis due to non-genetic effects, any variant that is also differentiated along this axis would have an overestimated effect size if the population structure is not well controlled in GWAS. Using GWAS summary statistics from a distant population should alleviate this issue, because the effect of stratification in the GWAS panel is independent from that of the testing populations for polygenic adaptation.

We first evaluated the impact of population stratification on height-associated variants ascertained from different GWAS panels that are available to us: the GIANT consortium, the UKB, and the BBJ datasets. Specifically, we examined the correlation between effect sizes estimated from each GWAS panel and the principal component loading on a PCA conducted in European populations (Supp Figure 1). We found that the effect sizes (beta) estimated in GIANT were highly correlated with the loading of the first two principal components of population structure (*rho* = 0.137, *p* = 2.49e-119 for PC1; *rho* = −0.018, *p* = 1.59e-4 for PC2) (Figure 1). Compared to the situation in GIANT, the correlations in UKB were much smaller, although still significant (*rho* = 0.017, *p* = 3.27e-3 for PC1; *rho* = −0.011, *p* = 7.08e-3 for PC2). In BBJ, the correlations were negligible (Figure 1). The lower degree of correlation observed in UKB could in principle be driven by selection, or alternatively the effect sizes on height estimated from UKB were not completely free from stratification. On the other hand, effect sizes from BBJ were independent of any population structure in Europeans. Therefore, we will conservatively use height-associated SNPs ascertained from BBJ as the set of SNPs used in primary analysis, although we also characterized the impact by ascertaining SNPs in different ascertainment panels for comparisons.

**Figure 1.**
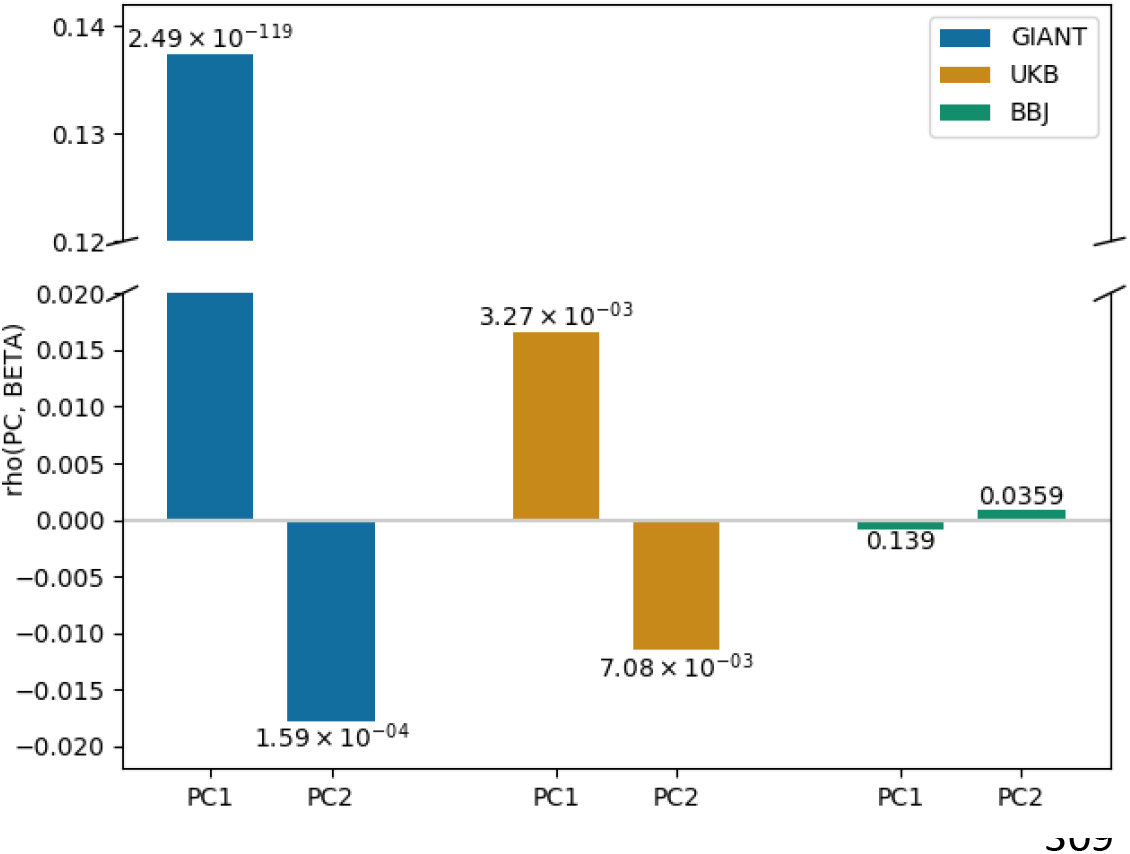
Evidence of stratification in GWAS summary statistics. Pearson Correlation coefficients of PC loadings and SNP effects from GIANT, UKB, and BBJ, across all SNPs. P values indicated on each bar are based on Jackknife standard errors (1000 blocks).

### Signals of polygenic adaptation in Sardinians

To evaluate the signal of polygenic adaptation in Sardinians, we calculated the average polygenic scores for Sardinians and four mainland European populations (CEU, TSI, GBR, and IBS) based on height-associated loci surpassing the genome-wide significant threshold (*P* < 5e-8) ascertained from GIANT, UKB, or BBJ. We then used Berg and Coop’s Q_x_ and conditional Z-score framework to evaluate the significance of differences in polygenic scores across populations. We found that across all GWAS panels the estimated polygenic height scores in Sardinians remain significantly lower than would be expected based on its genetic relatedness to European populations (Figure 2). However, the degree to which Sardinians were genetically shorter were more attenuated when using summary statistics derived from UKB and BBJ, relative to GIANT (Sardinians were 0.37, 0.35, and 0.94 units of s.d. shorter than CEU using polygenic scores computed from UKB, BBJ, and GIANT, respectively).

**Figure 2.**
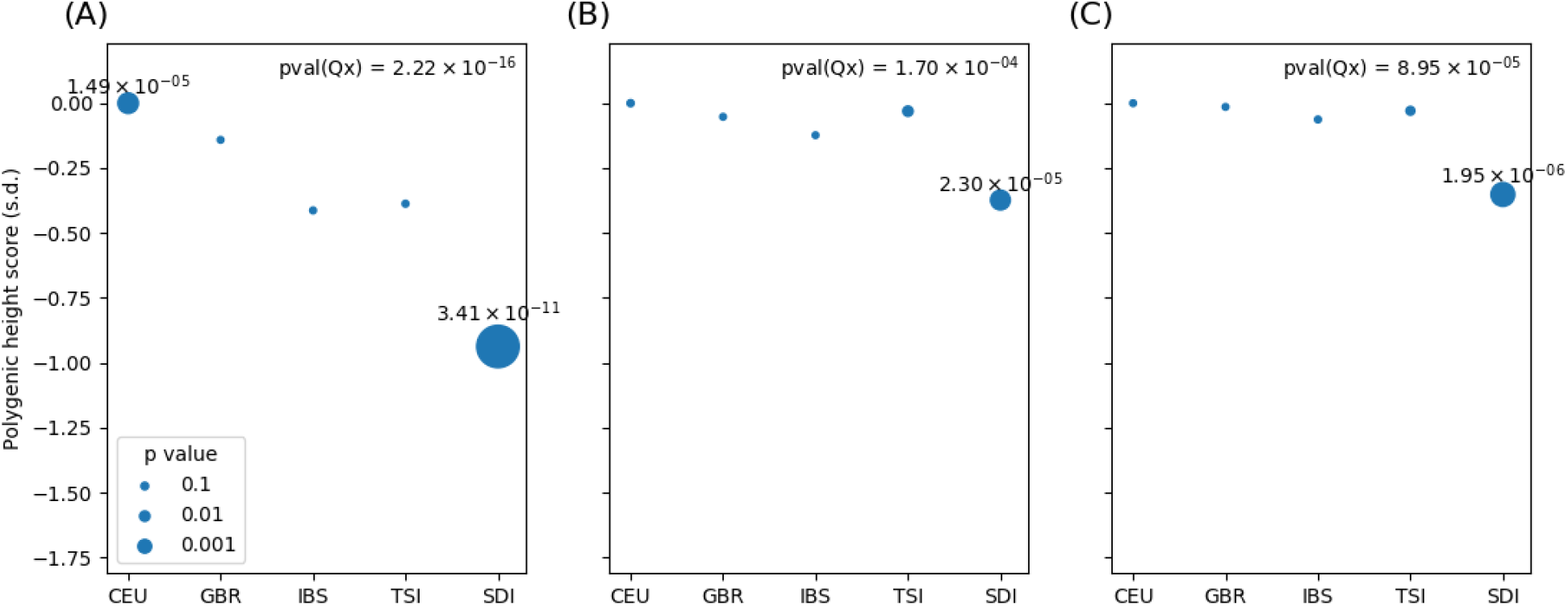
Excess variance tests in Sardinia. The polygenic score was constructed based on SNPs from (A) GIANT, (B) UKB, and (C) BBJ ascertained by picking genome-wide significant SNPs that are more than 500 kb apart from each other. The P values are represented by the size of each circle, and those lower than 0.01 are shown in the plot. SDI, Sardinians; IBS: Iberian Population in Spain, TSI: Toscani in Italia, GBR: British in England and Scotland, and CEU: Utah Residents with Northern and Western European Ancestry, are from 1000 Genome Project.

We focused on examining only variants surpassing genome-wide significance threshold to be better protected from uncorrected stratification, particularly when using summary statistics from European populations. But we also examined the effect of a different approach to identify height-associated variants used by previous reports (Berg et al., 2019; Sohail et al., 2019), namely by selecting the lowest p-value SNPs from approximately independent LD blocks across the genome. This resulted in approximately 1,700 variants using the GIANT or UKB GWAS panels, or 1,400 variants using the BBJ panel. By including more variants across the genome, most of which were not significantly associated with height, we observed that the polygenic height difference between Sardinians and the CEU population increased from 0.94 to 1.61, when using summary statistics from GIANT. More strikingly, the statistical evidence for adaptation became much stronger (P decreased from 3.41e-11 to 6.66e-16, Supp Figure 2). This is also observed, although to a lesser extent, when using summary statistics from UKB (Supp Figure 2). In contrast, when using summary statistics from BBJ, this ascertainment scheme did not significantly exacerbate the difference in polygenic height scores between Sardinia and CEU, and actually decreased the statistical evidence of adaptation. In fact, ascertaining variants from BBJ summary statistics provided the strongest signal-to-noise ratio in the per-variant contribution of population differences (14.22 vs. 6.64 when using UKB; Table 1). When we stratified variants based on whether the variant surpassed the genome-wide significance threshold, we found the per-variant contributions of polygenic height score differences between CEU and Sardinians due to sub-threshold variants in the GIANT summary statistics (7.15e-4 s.d. difference per allele) to be as large as that due to genome-wide significant variants in the UKB or BBJ summary statistics (4.22e-4 and 7.18e-4 s.d. difference per allele), consistent with enrichment of uncontrolled stratification (Table 1). On the other hand, the contributions due to sub-threshold variants in BBJ summary statistics were an order of magnitude less than that of the genome-wide significant variants (Table 1). Taken together, these results suggest that the exaggerated signature of polygenic adaptation using GIANT was at least partly due to the practice of ascertain SNPs in approximate linkage equilibrium blocks, which enriched for loci that escaped statistical control of stratification. A better analytical practice would be to analyze a set of independent variants ranked by p-values such that true height-associated variants will be highly enriched. This also attested that summary statistics from BBJ is free from stratification in the test (European) populations, as sub-threshold SNPs added noise to the analysis.

**Table 1.** Per-variant polygenic height score differences between CEU and Sardinians

|  | Independent LD block (# variants) | Genome-wide significant variants | Nonsignificant variants | Ratio between significant and nonsignificant variants |
| --- | --- | --- | --- | --- |
| GIANT | 9.46e-04 (1702) | 1.54e-03 (476) | 7.15e-04 (1226) | 2.15 |
| UKB | 2.34e-04 (1703) | 4.22e-04 (812) | 6.36e-05 (891) | 6.64 |
| BBJ | 2.26e-04 (1444) | 7.18e-04 (379) | 5.05e-05 (1065) | 14.22 |

The interpretation of the conditional Z-score results (Figure 2) implicitly assumes that all other populations tested in this framework are evolving neutrally. Our observation could also be explained if height-associated loci are evolving neutrally in Sardinia, but collectively increasing the genetic height in all mainland European populations. To further investigate if selection is acting on height-associated loci in Sardinia, we compared the trajectory of polygenic height scores in Sardinia to that from the GBR population. Using the proportion-of-lineages estimator from Edge and Coop (Edge & Coop, 2019), the most powerful of the three estimators introduced in terms of detecting adaptation, we observed that the population-mean polygenic score for height has been decreasing in Sardinia, while it mainly stayed constant in GBR, in the past ~10 thousand years (ky) (Figure 3). We tested if whether the decreasing trend in Sardinia was significant relative to GBR for two time points: 20 ky and 10 ky before present, as these were approximately the time with first evidence of human inhabitation on Sardinia (Calò et al., 2008; Vona, 1997) and the Neolithic period and divergence between present day Sardinians and mainland Europeans (Chiang et al., 2018), respectively. We found that the mean polygenic height score in Sardinia was significantly decreasing from that in GBR since at least 10 ky ago (kya) (p = 0.0123). The trend was not significant between 20 ky and 10 ky ago (p = 0.5307). We also observed qualitatively similar trend of decrease in polygenic height score in Sardinia relative to GBR using the other two estimators presented in Edge and Coop (Supp Figure 3), however, these estimators are known to be much more variable and thus have much less power to detect selection (Edge & Coop, 2019). When using height-associated loci ascertained from UKB, we observed similar results (Supp Figure 4; *p* = 0.0321 and 0.5071 for differences between 10 kya to present and 20 kya to 10 kya, respectively).

**Figure 3.**
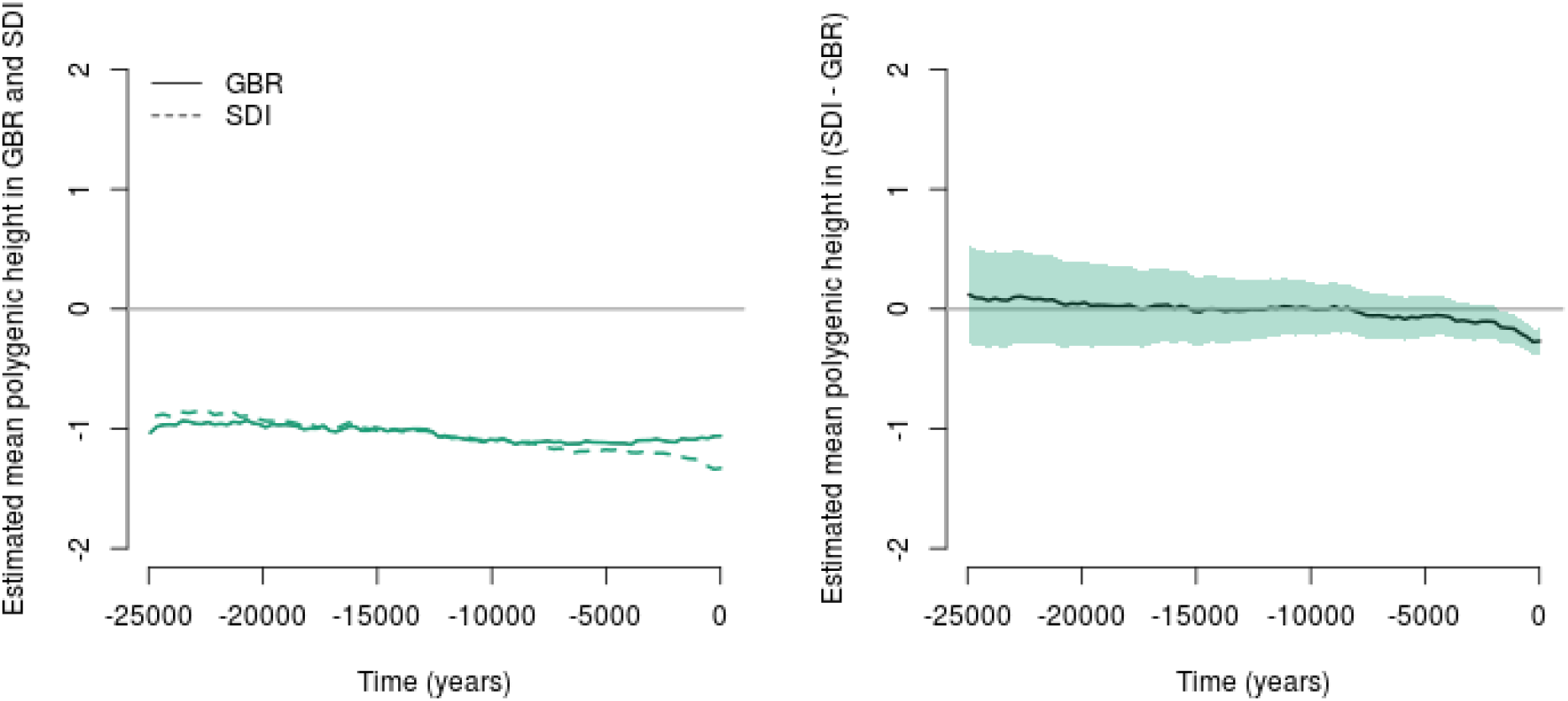
The trajectory of mean polygenic height scores in British (GBR) and Sardinian (SDI) populations over the past 25 ky. The past polygenic scores are estimated by the proportion-of-lineages estimators from Edge and Coop, using height loci and effect sizes ascertained from BBJ. The left panel shows the mean polygenic scores in the GBR and SDI using the proportion-of-lineages estimator. The right panel shows the difference between GBR and SDI in the mean polygenic scores using the estimator. Shaded areas denote the 95% confidence interval. The mean polygenic height score in Sardinia was significantly decreasing from that in GBR since at least 10 kya (p = 0.0123), while not significant between 20 kya and 10 kya (p = 0.5307).

### Signals of polygenic adaptation in mainland Europeans

We then focused on evaluating whether there is a signal of polygenic adaptation in mainland Europeans, the population originally indicated (Berg & Coop, 2014; Field et al., 2016; Robinson et al., 2015; Turchin et al., 2012) but recently challenged (Berg et al., 2019; Sohail et al., 2019). Consistent with the recent reports, using Berg and Coop’s Q_x_ and conditional Z-score, we observed clear signal of adaptation when ascertaining based on summary statistics from GIANT, but not UKB or BBJ (Supp Figure 5).

Any signal of adaptation in the mainland Europe, if it exists, is undoubtedly weaker than that in Sardinia. Therefore, we further assessed if the lack of signal in the analysis based on variants ascertained from BBJ could be due to loss of power in the Q_x_ and conditional Z-score. We explored two potential causes. First, we examined the frequency distribution of BBJ height variants in mainland Europeans. We found that there is a shift towards rarer alleles in BBJ. The mean MAF in Europe is 0.227 and 0.258 for BBJ-ascertained and UKB-ascertained variants, respectively (Supp Figure 6). Although the majority of BBJ-ascertained variants had relatively high frequency in European (~350 SNPs, or 74.32% of all BBJ-ascertained variants, with MAF > 0.1), the rarer variants also tend to be less associated with height in UKB (Supp Figure 7). For example, of the 108 variants with MAF < 0.1 in UKB, 47.22% of them are not associated with height (p-value > 0.05), suggesting possibly that differences in LD between UKB and BBJ dissociated the causal SNP from the proxy we selected, or that the effect on height is specific to BBJ. However, despite the additional noises from potentially selecting the “wrong” variants for analysis, rerunning Q_x_ test (Berg & Coop, 2014) restricting to BBJ variants with MAF > 0.1 in mainland Europeans or with P-value < 5e-8 in UKB did not qualitatively change the results (Supp Figure 8).

Secondly, the Conditional Z-test relies on testing a particular population while conditioning on relationship to all other populations in the dataset, assuming all other populations are neutrally evolving. If, for example, a very closely related population (e.g., GBR to CEU) is included in the reference population and assumed to be evolving neutrally, this could decrease the power to detect adaptation using this framework. Therefore, we conducted a simple t-test of frequency difference of height-increasing alleles (Turchin et al., 2012) at genome-wide significant variants between Northern Europeans (grouping CEU and GBR) and Southern European (grouping IBS and TSI). Using height-associated loci ascertained from GIANT, we observe clear difference in frequency between the Northern and Southern Europeans (height-increasing alleles are on average 1.12% higher in Northern Europeans, *p* = 5.24e-8). On the other hand, using height-associated loci ascertained from either UKB or BBJ, we found much more attenuated signals that are not significant (0.011% and 0.20%, *p* = 0.931 and 0.313 in UKB and BBJ, respectively), even if we applied more permissive p-value thresholds on both populations (Supp Table 1).

Because the sample sizes in mainland European populations from 1000 Genomes are relatively small (N = 190 for Northern Europeans, 214 for Southern Europeans), the estimate of allele frequency may suffer from sampling errors that could mask the allele frequency difference between populations. We found it difficult to obtain precise estimates of allele frequency ideally based on thousands of individuals, particularly for Southern Europeans on the mainland. To partly overcome this challenge, we downloaded variant frequency data from 125,748 exomes from gnomAD v2.1 (Genome Aggregation Database) (Karczewski et al., 2019), which contained 21,111 samples categorized as North-Western Europeans (NWE), and 5,752 samples categorized as Southern Europeans (SEU). We restricted our analysis to only independent exome variants which have frequency estimated from more than 10,000 alleles in both SEU and NEW and tested in the GWAS panel. We found no significant differences in frequency again after adjusting for the number of tests performed (Supp Table 2). However, note that the lack of signal could be at least partly attributed to the relatively low LD between the tagging exomic variants and the nearby top variants in the GWAS summary statistics (Supp Figure 9), as well as the constraints due to purifying selection that the exonic variants are expected to also experience.

While the allele frequency-based approaches deployed so far would be subjected to a loss of power due to incompleteness in the frequency estimates on the exact variants in large cohorts, haplotype-based analysis may be more sensitive for detecting adaptation. We thus evaluated the signal of polygenic adaptation in 3,195 individuals from UK10K dataset using the Singleton Density Score polarized to the height-increasing allele (tSDS), which should be most powered to detect changes over the last 2,000-3,000 years (Field et al., 2016). Focusing on independent variants surpassing genome-wide significance in either UKB or BBJ, we found that tSDS for height-increasing alleles are significantly elevated (*p* = 1e-5 and 4.8e-4 when ascertaining from UKB and BBJ, respectively; Figure 4). This suggests that the height-increasing alleles, compared to the height-decreasing alleles at the same genomic site, is more likely to be found on the longer haplotype, consistent with positive selection in the recent past. The result using height-associated variants ascertained from BBJ is particularly encouraging, as the patterns of variation at these loci appear unaffected by the major axis of stratification and height differentiation within Europe (Figure 1). While our conclusion is limited to the British population for which large-scale genome-wide sequencing data is available, taken together our results do imply that outside of the Sardinian population, height differences in at least some populations in mainland Europe may be driven by polygenic selection. Future sequencing data or more precise genome-wide frequency estimates may further characterize this observation.

**Figure 4.**
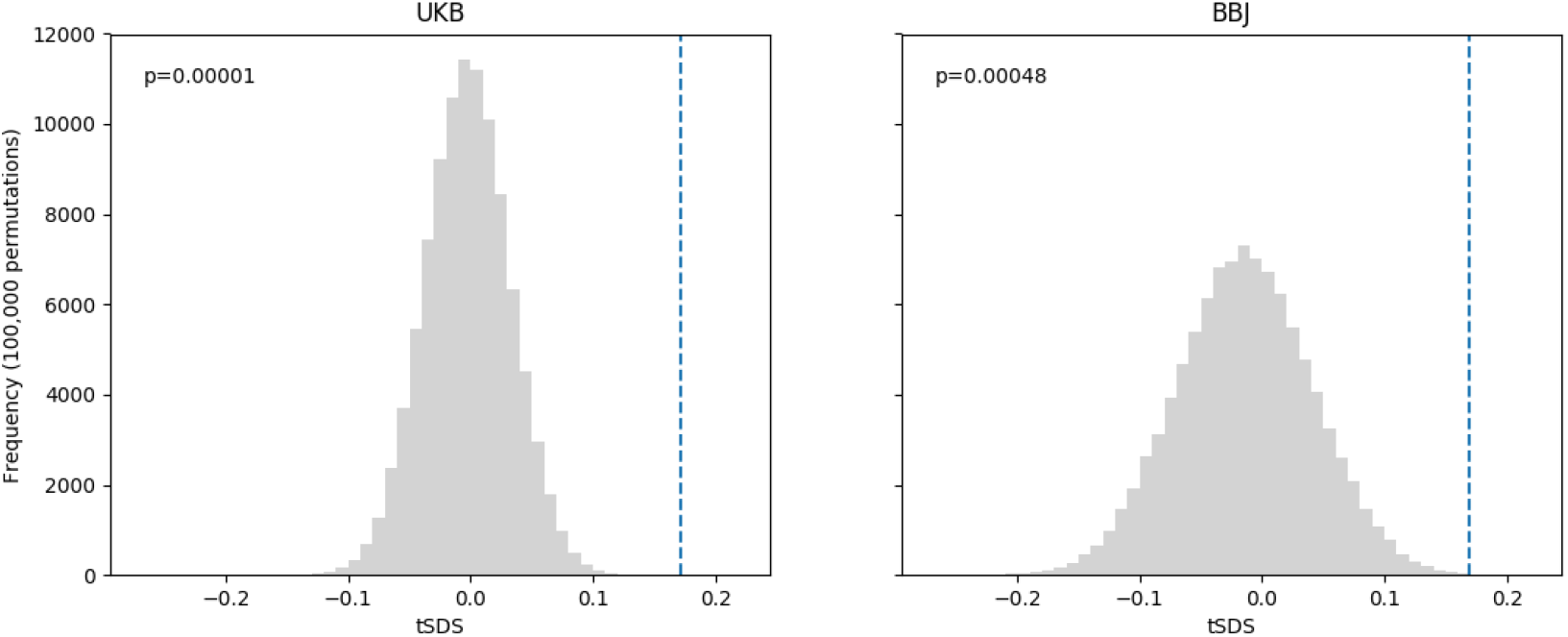
The average of tSDS in height-associated SNPs in UKB and BBJ. The histogram is the null distribution of average tSDS from 100,000 permutations. The dashed line indicates the observed average of tSDS.

## Discussion

Using a set of height-associated alleles ascertained from a GWAS panel that is not impacted by population structure in Europe (Figure 1), we have demonstrated in this study that height alleles appear to be under selection in some European populations. Our observation among the Sardinians is qualitatively consistent with previous report in this population (Zoledziewska et al., 2015), although the signal is more attenuated (Figure 2). Moreover, using the recently developed method to infer trajectory of polygenic height scores, we showed that the strongest feature in our analysis is driven by a decrease in polygenic height scores in the Sardinians. In mainland Europe, we could not detect a signature of polygenic adaptation using frequency-based methods (Supp Figures 5 and 8; Supp Tables 1 and 2), consistent with previous suggestions (Berg et al., 2019; Sohail et al., 2019) that selection signals, if any, would be much more attenuated. However, we did find a robust signal of recent selection for height-increasing alleles in an independent British population using tSDS (Figure 4).

A major consideration in detecting signals of polygenic selection is to examine the causal SNPs for the trait of interest. As we found in Table 1, variants that escaped state-of-the-art statistical controls for stratification in GWAS could be enriched among those ascertained from approximately independent LD blocks. Focusing only on genome-wide significant variants can help alleviate the concerns of population stratification. (Although the effect sizes may still reflect some residual stratification in a polygenic score style of analysis.) On the other hand, trait-associated variants found in GWAS using genotyping data are only proxies for causal variation. Differences in LD across population could thus lead to a decrease in the accuracy of polygenic score prediction and the lowered power in detecting polygenic selection. Fine-mapping could help identify the causal or the best tagging variants at associated loci. We elected not to conduct a fine-mapping analysis because currently the largest GWAS datasets are in Europeans and East Asians; fine-mapping approaches using Europeans may cause residual stratification to seep into the ascertainment scheme. As larger non-European GWAS datasets become available, fine-mapping studies outside of Europe may provide better set of causal alleles associated with height to address the question of adaptation in Europe.

A second consideration is that the lack of signals in mainland Europeans using direct comparison of allele frequencies might be partly due to the small sample sizes in publicly available, geographically indexed, whole genome sequences. Because only a subtle allele frequency shift would be expected in mainland Europe, the imprecision in allele frequency estimates can mask the signal of adaptation (subtle coordinated allele frequency shifts between populations). We note that Berg and Coop’s framework of excess variance test and conditional Z-scores (Berg & Coop, 2014) could also possibly be impacted by this issue with precision of allele frequency estimates.

Concerns of both the imprecise ascertainment of causal allele and of allele frequency estimates could be partly overcome by large-scale haplotype-based methods. Using height-associated loci ascertained from BBJ, where stratification should no longer be an issue, we observed a robust signal of recent selection for height-increasing alleles using tSDS, which was calculated on a sample size of > 3,000 UK individuals (Field et al., 2016) (Figure 4). In contrast, our polygenic score trajectory analysis did not show any recent increase in the 91 GBR individuals (Figure 3). This analysis is a hybrid approach based on an inferred ancestral recombination graph that combines both haplotype and genotype information, so the same shortcomings in frequency-based approach could partly explain the null finding. Moreover, computationally this approach is currently limited to smaller sample sizes, which may also limit our resolution in the recent past. Therefore, future scalable inference on genome-wide genealogies from an independent Northern European population may help address this discrepancy.

The signal of natural selection favoring shorter stature in the Sardinians is very strong, regardless of whether we used loci ascertained from UKB or BBJ. The trajectory of polygenic height scores in the Sardinians decreased significantly from that in GBR during the past 10,000 years further supported this conclusion. Taking into account recent evidence of selection for shorter stature in the island of Flores, these observations might suggest a general impact due to the island effect, akin to what has been observed in some island mammals, who became adaptively smaller relative to their mainland counterparts (Millien, 2006; model 1 in Figure 5). However, as the power of polygenic score trajectory to detect between-population differences decreases going further back in time (Edge & Coop, 2019), we note that we cannot definitively infer that adaptation towards shorter height occurred on the Sardinia island starting around 10,000 years ago. It is possible that adaptation occurred in the ancestors of modern Sardinians, long before the Sardinia island was populated. Because of the relative isolation since the founding of Sardinia (Chiang et al., 2018), the Sardinian population exhibit the strongest effect among European populations today (model 2 in Figure 5). This may be consistent with the recent observation that Neolithic European populations are shorter than both their predecessors and their successors in Europe in both genetic and skeletal stature (Cox, Ruff, Maier, & Mathieson, 2019); Sardinians retained the largest amount of Neolithic ancestry among a number of extant European populations tested (Chiang et al., 2018; Haak et al., 2015). Furthermore, the two models are not mutually exclusive, and may be acting along with non-additive components of height variation (Joshi et al., 2015; Zoledziewska et al., 2015) to lead to the large difference in height observed between Sardinians and mainland Europeans. When considering Europe at large, our tSDS findings in the British population may be consistent with previous suggestions that the post-Neolithic Eurasian steppe populations may have been selected for increased height (Martiniano et al., 2017), and that admixture of these populations, in different proportions in mainland Europe, provided the tSDS signal and contributed to the pattern of height variation across Europe. If such an event occurred, it seems that it would have been independent of a selection for shorter height in Sardinians or their ancestors, since we observed a monotonic decreasing trend in polygenic height in Sardinians. Further explorations to differentiate these two models will likely rely on examining a large number of ancient specimen from Sardinia (Marcus et al., 2019), as well as studying other isolated island populations across the world.

Taken together, while the timing and geographical location of selection for height (or alternatively, for a set of traits correlated with height) remain elusive, it seems evident that human height differences in Europe had been driven by selection in at least some instances. Multiple episodes of adaptation may have occurred and influenced the height of past populations. Signatures of these adaptive events may have stemmed from outside of Sardinia but today they are much more obscured, or even changed direction, due to recent population migrations and admixture. Furthermore, while much of the literature characterizing polygenic scores stemming from GWAS summary statistics have focused on its poor transferability between populations (Martin et al., 2019), our results here demonstrate that such a polygenic score estimator is also unbiased and can greatly help addressing important population genetic questions.

**Figure 5.**
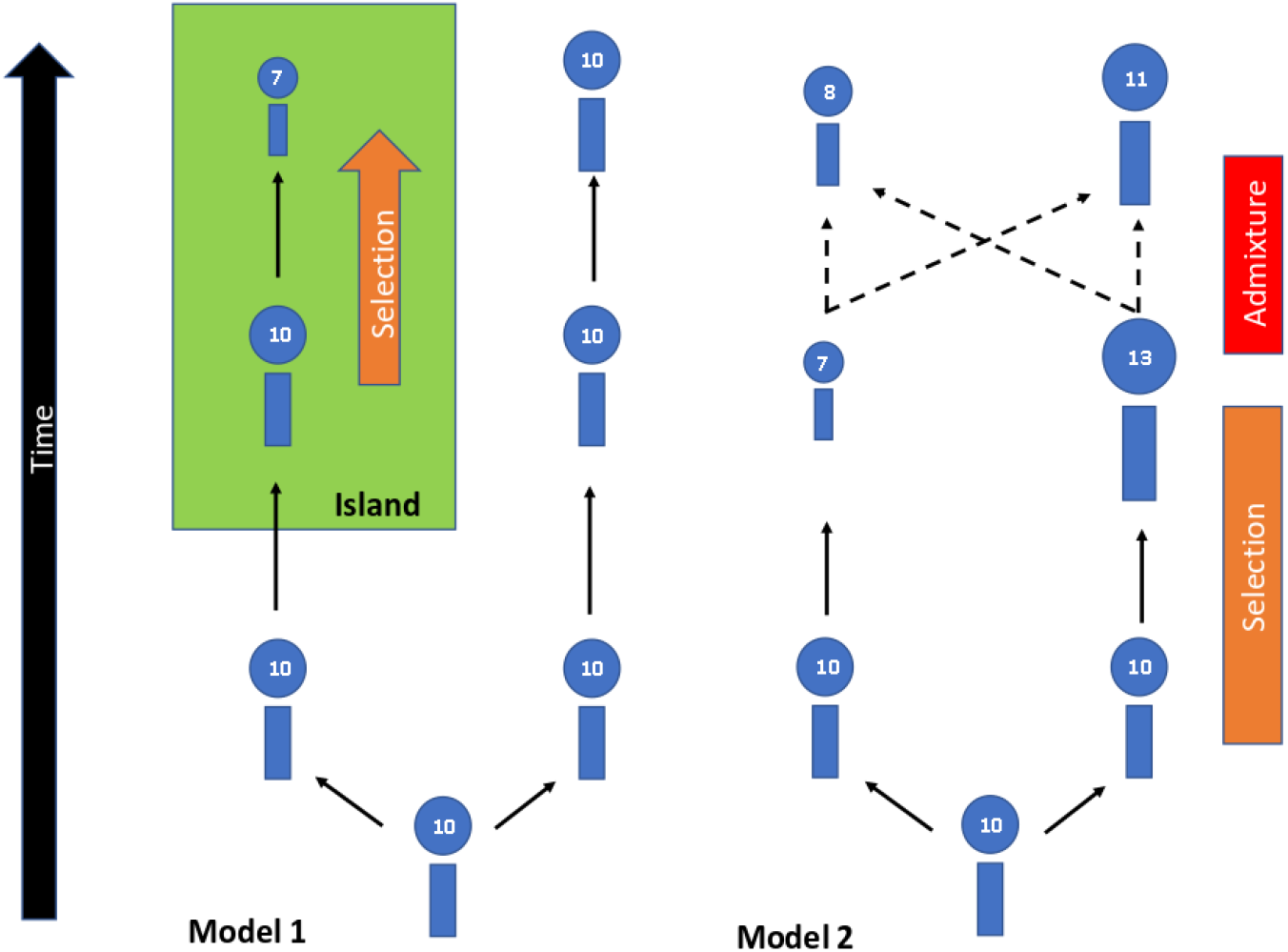
Possible evolutionary models for human height. Each figure represents a modern or ancestral population, with a pseudo score for height labeled in the circle. Model 1 (left) illustrates the island effect, in which an island population, possibly like the one in Sardinia, were selected for shorter stature upon arriving at the island. The non-island population were assumed to be not under selection. Model 2 (right) illustrates an alternative model, in which selection occurred much more anciently such that differentiation between populations has occurred. It is unclear the direction of selection in this scenario, and may be in opposite directions in different populations due to differential interaction with the geography or environment. At this point, height no longer needs to be selected anymore, but subsequent migration between populations establishes the pattern of height variability. In both cases, adaptation has occurred and can be detected by the framework utilized in this paper, although we cannot infer the timing and the location of selection. Also note that instead of height, a trait or collection of traits proxied by height could be under selection, although we focused on height in this model.

## Supplemental Data

Supplemental Data include 9 figures and 2 tables.

## Acknowledgments

We gratefully thank Graham Coop, Michael D. Edge, and Michael C. Turchin for helpful comments and discussions. This work is supported by Start-up funds provided by the Center for Genetic Epidemiology at Keck School of Medicine USC (to C.W.K.C.).

## Web Resources

GIANT summary association statistics, https://portals.broadinstitute.org/collaboration/giant/images/0/01/GIANT_HEIGHT_Wood_et_al_2014_publicrelease_HapMapCeuFreq.txt.gz

UKB summary association statistics, https://storage.googleapis.com/ukbb-robert/height_ukb_giant/robert1/50.imputed_v3.results.both_sexes.tsv.gz

Genetic maps generated from the 1000 Genome phased OMNI data, http://ftp.1000genomes.ebi.ac.uk/vol1/ftp/technical/working/20130507_omni_recombination_rates/

Human ancestral genome, ftp://ftp.1000genomes.ebi.ac.uk/vol1/ftp/phase1/analysis_results/supporting/ancestral_alignments/

Genomic mask file, ftp://ftp.1000genomes.ebi.ac.uk/vol1/ftp/release/20130502/supporting/accessible_genome_masks/

SDS from UK10K, http://web.stanford.edu/group/pritchardlab/UK10K-SDS-values.zip

PLINK version 1.9, www.cog-genomics.org/plink/1.9/

Eigensoft version 7.2.1, https://github.com/DReichLab/EIG/archive/v7.2.1.tar.gz

Eagle version 2.4.1, https://data.broadinstitute.org/alkesgroup/Eagle/

RELATE version 1.0.8, https://myersgroup.github.io/relate/

Code for Q_x_ test, https://github.com/jjberg2/PolygenicAdaptationCode

Code for polygenic score trajectory, https://github.com/mdedge/rhps_coalescent

## Reference

1000 Genomes Project Consortium, Auton, A., Brooks, L. D., Durbin, R. M., Garrison, E. P., Kang, H. M., … Abecasis, G. R. (2015). A global reference for human genetic variation. Nature, 526(7571), 68–74. https://doi.org/10.1038/nature15393

Akiyama, M., Ishigaki, K., Sakate, S., Momozawa, Y., Horikoshi, M., Hirata, M., & Kamatani, Y. (n.d.). Characterizing rare and low-frequency height-associated variants in the Japanese population. Nature Communications.

Bentham, J., Di Cesare, M., Stevens, G. A., Zhou, B., Bixby, H., Cowan, M., … Cisneros, J. Z. (2016). A century of trends in adult human height. ELife, 5(2016 JULY). https://doi.org/10.7554/eLife.13410.001

Berg, J. J., & Coop, G. (2014). A Population Genetic Signal of Polygenic Adaptation. PLoS Genetics, 10(8), e1004412. https://doi.org/10.1371/journal.pgen.1004412

Berg, J. J., Harpak, A., Sinnott-Armstrong, N., Joergensen, A. M., Mostafavi, H., Field, Y., … Coop, G. (2019). Reduced signal for polygenic adaptation of height in UK Biobank. ELife, 8. https://doi.org/10.7554/eLife.39725

Berisa, T., & Pickrell, J. K. (2016). Approximately independent linkage disequilibrium blocks in human populations. Bioinformatics (Oxford, England), 32(2), 283–285. https://doi.org/10.1093/bioinformatics/btv546

Calò, C., Melis, A., Vona, G., & Piras, I. (2008). Review Synthetic Article: Sardinian Population (Italy): a Genetic Review. International Journal of Modern Anthropology, 1(1), 39–64. https://doi.org/10.4314/ijma.v1i1.60356

Chang, C. C., Chow, C. C., Tellier, L. C., Vattikuti, S., Purcell, S. M., & Lee, J. J. (2015). Second-generation PLINK: rising to the challenge of larger and richer datasets. GigaScience, 4(1), 7. https://doi.org/10.1186/s13742-015-0047-8

Chiang, C. W. K., Marcus, J. H., Sidore, C., Biddanda, A., Al-Asadi, H., Zoledziewska, M., … Novembre, J. (2018). Genomic history of the Sardinian population. Nature Genetics, 50(10), 1426–1434. https://doi.org/10.1038/s41588-018-0215-8

Cox, S. L., Ruff, C. B., Maier, R. M., & Mathieson, I. (2019). Genetic contributions to variation in human stature in prehistoric Europe. BioRxiv, 690545. https://doi.org/10.1101/690545

Edge, M. D., & Coop, G. (2019). Reconstructing the History of Polygenic Scores Using Coalescent Trees. Genetics, 211(1), 235–262. https://doi.org/10.1534/genetics.118.301687

Field, Y., Boyle, E. A., Telis, N., Gao, Z., Gaulton, K. J., Golan, D., … Pritchard, J. K. (2016). Detection of human adaptation during the past 2000 years. Science, 354(6313), 760–764. https://doi.org/10.1126/SCIENCE.AAG0776

Grasgruber, P., Cacek, J., Kalina, T., & Sebera, M. (2014). The role of nutrition and genetics as key determinants of the positive height trend. Economics & Human Biology, 15, 81–100. https://doi.org/10.1016/J.EHB.2014.07.002

Guo, J., Wu, Y., Zhu, Z., Zheng, Z., Trzaskowski, M., Zeng, J., … Yang, J. (2018). Global genetic differentiation of complex traits shaped by natural selection in humans. Nature Communications, 9(1), 1865. https://doi.org/10.1038/s41467-018-04191-y

Haak, W., Lazaridis, I., Patterson, N., Rohland, N., Mallick, S., Llamas, B., … Reich, D. (2015). Massive migration from the steppe was a source for Indo-European languages in Europe. Nature, 522(7555), 207–211. https://doi.org/10.1038/nature14317

Haworth, S., Mitchell, R., Corbin, L., Wade, K. H., Dudding, T., Budu-Aggrey, A., … Timpson, N. J. (2019). Apparent latent structure within the UK Biobank sample has implications for epidemiological analysis. Nature Communications 2019 10:1, 10(1), 333. https://doi.org/10.1038/s41467-018-08219-1

Hirata, M., Kamatani, Y., Nagai, A., Kiyohara, Y., Ninomiya, T., Tamakoshi, A., … Yoshiyama, T. (2017). Cross-sectional analysis of BioBank Japan clinical data: A large cohort of 200,000 patients with 47 common diseases. Journal of Epidemiology, 27(3), S9–S21. https://doi.org/10.1016/j.je.2016.12.003

Joshi, P. K., Esko, T., Mattsson, H., Eklund, N., Gandin, I., Nutile, T., … Wilson, J. F. (2015). Directional dominance on stature and cognition in diverse human populations. Nature, 523(7561), 459–462. https://doi.org/10.1038/nature14618

Karczewski, K. J., Francioli, L. C., Tiao, G., Cummings, B. B., Alföldi, J., Wang, Q., … MacArthur, D. G. (2019). Variation across 141,456 human exomes and genomes reveals the spectrum of loss-of-function intolerance across human protein-coding genes. BioRxiv, 531210. https://doi.org/10.1101/531210

Kerminen, S., Martin, A. R., Koskela, J., Ruotsalainen, S. E., Havulinna, A. S., Surakka, I., … Pirinen, M. (2019). Geographic Variation and Bias in the Polygenic Scores of Complex Diseases and Traits in Finland. The American Journal of Human Genetics, 104(6), 1169–1181. https://doi.org/10.1016/j.ajhg.2019.05.001

Liu, X., Loh, P.-R., O’Connor, L. J., Gazal, S., Schoech, A., Maier, R. M., … Price, A. L. (2018). Quantification of genetic components of population differentiation in UK Biobank traits reveals signals of polygenic selection. BioRxiv, 357483. https://doi.org/10.1101/357483

Locke, A. E., Steinberg, K. M., Chiang, C. W. K., Service, S. K., Havulinna, A. S., Stell, L., … Freimer, N. B. (2019). Exome sequencing of Finnish isolates enhances rare-variant association power. Nature, 1–6. https://doi.org/10.1038/s41586-019-1457-z

Loh, P.-R., Tucker, G., Bulik-Sullivan, B. K., Vilhjálmsson, B. J., Finucane, H. K., Salem, R. M., … Price, A. L. (2015). Efficient Bayesian mixed-model analysis increases association power in large cohorts. Nature Genetics, 47(3), 284–290. https://doi.org/10.1038/ng.3190

Loh, P. R., Danecek, P., Palamara, P. F., Fuchsberger, C., Reshef, Y. A., Finucane, H. K., … Price, A. L. (2016). Reference-based phasing using the Haplotype Reference Consortium panel. Nature Genetics, 48(11), 1443–1448. https://doi.org/10.1038/ng.3679

Marcus, J. H., Posth, C., Ringbauer, H., Lai, L., Skeates, R., Sidore, C., … Novembre, J. (2019). Population history from the Neolithic to present on the Mediterranean island of Sardinia: An ancient DNA perspective. BioRxiv, 583104. https://doi.org/10.1101/583104

Martin, A. R., Kanai, M., Kamatani, Y., Okada, Y., Neale, B. M., & Daly, M. J. (2019). Clinical use of current polygenic risk scores may exacerbate health disparities. Nature Genetics, 51(4), 584–591. https://doi.org/10.1038/s41588-019-0379-x

Martiniano, R., Cassidy, L. M., O’Maolduin, R., McLaughlin, R., Silva, N. M., Manco, L., … Bradley, D. G. (2017). The population genomics of archaeological transition in west Iberia: Investigation of ancient substructure using imputation and haplotype-based methods. PLoS Genet, 13(7), e1006852. https://doi.org/10.1371/journal.pgen.1006852

McVicker, G., Gordon, D., Davis, C., & Green, P. (2009). Widespread Genomic Signatures of Natural Selection in Hominid Evolution. PLoS Genetics, 5(5), e1000471. https://doi.org/10.1371/journal.pgen.1000471

Millien, V. (2006). Morphological Evolution Is Accelerated among Island Mammals. PLoS Biology, 4(10), e321. https://doi.org/10.1371/journal.pbio.0040321

Nagai, A., Hirata, M., Kamatani, Y., Muto, K., Matsuda, K., Kiyohara, Y., … Kubo, M. (2017). Overview of the BioBank Japan Project: Study design and profile. https://doi.org/10.1016/j.je.2016.12.005

Price, A. L., Weale, M. E., Patterson, N., Myers, S. R., Need, A. C., Shianna, K. V., … Reich, D. (2008). Long-Range LD Can Confound Genome Scans in Admixed Populations. American Journal of Human Genetics. https://doi.org/10.1016/j.ajhg.2008.06.005

Robinson, M. R., Hemani, G., Medina-Gomez, C., Mezzavilla, M., Esko, T., Shakhbazov, K., … Visscher, P. M. (2015). Population genetic differentiation of height and body mass index across Europe. Nature GeNetics VOLUME, 47, 34. https://doi.org/10.1038/ng.3401

Sohail, M., Maier, R. M., Ganna, A., Bloemendal, A., Martin, A. R., Turchin, M. C., … Sunyaev, S. R. (2019). Polygenic adaptation on height is overestimated due to uncorrected stratification in genome-wide association studies. ELife, 8. https://doi.org/10.7554/eLife.39702

Speidel, L., Forest, M., Shi, S., & Myers, S. R. (2019). A method for genome-wide genealogy estimation for thousands of samples. Nature Genetics, 51(9), 1321–1329. https://doi.org/10.1038/s41588-019-0484-x

Tucci, S., Vohr, S. H., McCoy, R. C., Vernot, B., Robinson, M. R., Barbieri, C., … Green, R. E. (2018). Evolutionary history and adaptation of a human pygmy population of Flores Island, Indonesia. Science (New York, N.Y.), 361(6401), 511–516. https://doi.org/10.1126/science.aar8486

Turchin, M. C., Chiang, C. W., Palmer, C. D., Sankararaman, S., Reich, D., & Hirschhorn, J. N. (2012). Evidence of widespread selection on standing variation in Europe at height-associated SNPs. Nature Genetics, 44(9), 1015–1019. https://doi.org/10.1038/ng.2368

Vona, G. (1997). The peopling of Sardinia (Italy): history and effects. International Journal of Anthropology, 12(1), 71–87. https://doi.org/10.1007/BF02447890

Wood, A. R., Esko, T., Yang, J., Vedantam, S., Pers, T. H., Gustafsson, S., … Frayling, T. M. (2014). Defining the role of common variation in the genomic and biological architecture of adult human height. Nature Genetics, 46(11), 1173–1186. https://doi.org/10.1038/ng.3097

Zoledziewska, M., Sidore, C., Chiang, C. W. K., Sanna, S., Mulas, A., Steri, M., … Cucca, F. (2015). Height-reducing variants and selection for short stature in Sardinia. Nature Genetics, 47(11), 1352–1356. https://doi.org/10.1038/ng.3403

