## Supplemental Material for "Evidence of polygenic adaptation at height-associated loci in mainland Europeans and Sardinians"

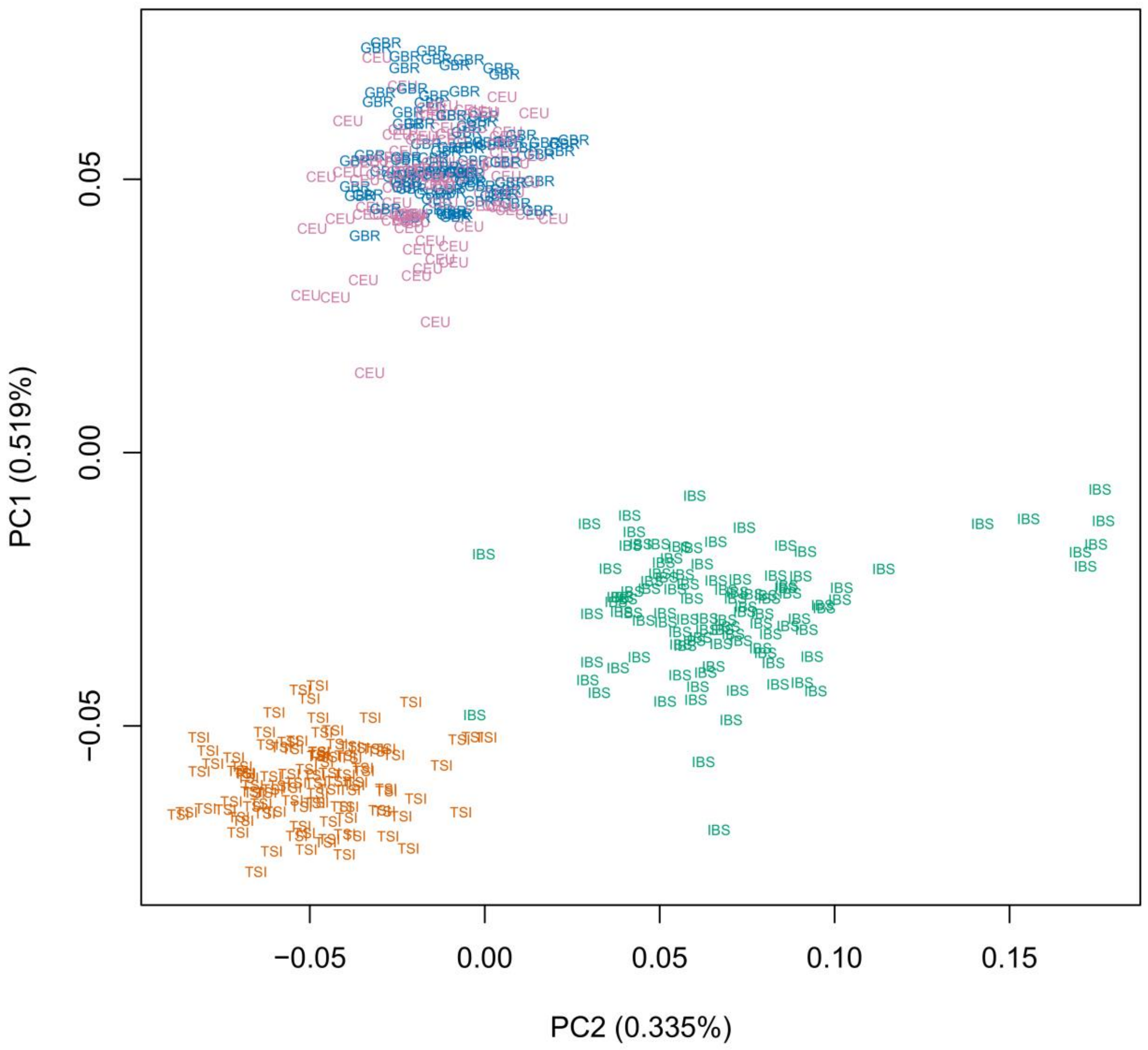


Supp Figure 1. The first two principal components in PCA on mainland Europeans from 1000 Genome. IBS: Iberian Population in Spain, TSI: Toscani in Italia, GBR: British in England and Scotland, and CEU: Utah Residents with Northern and Western European Ancestry.

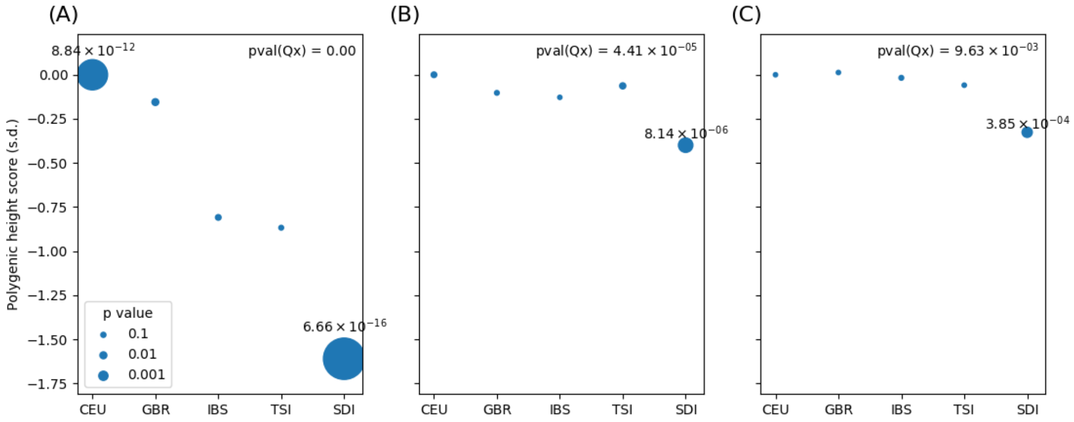


Supp Figure 2. Excess variance tests in Sardinia. The polygenic score was constructed based on SNPs from (A) GIANT, (B) UKB, and (C) BBJ ascertained by picking the SNP with the lowest *P* value in each approximately independent LD block. The *P* values are represented by the size of each circle, and those lower than 0.01 are shown in the plot. SDI, Sardinians; IBS: Iberian Population in Spain, TSI: Toscani in Italia, GBR: British in England and Scotland, and CEU: Utah Residents with Northern and Western European Ancestry, are from 1000 Genome Project.

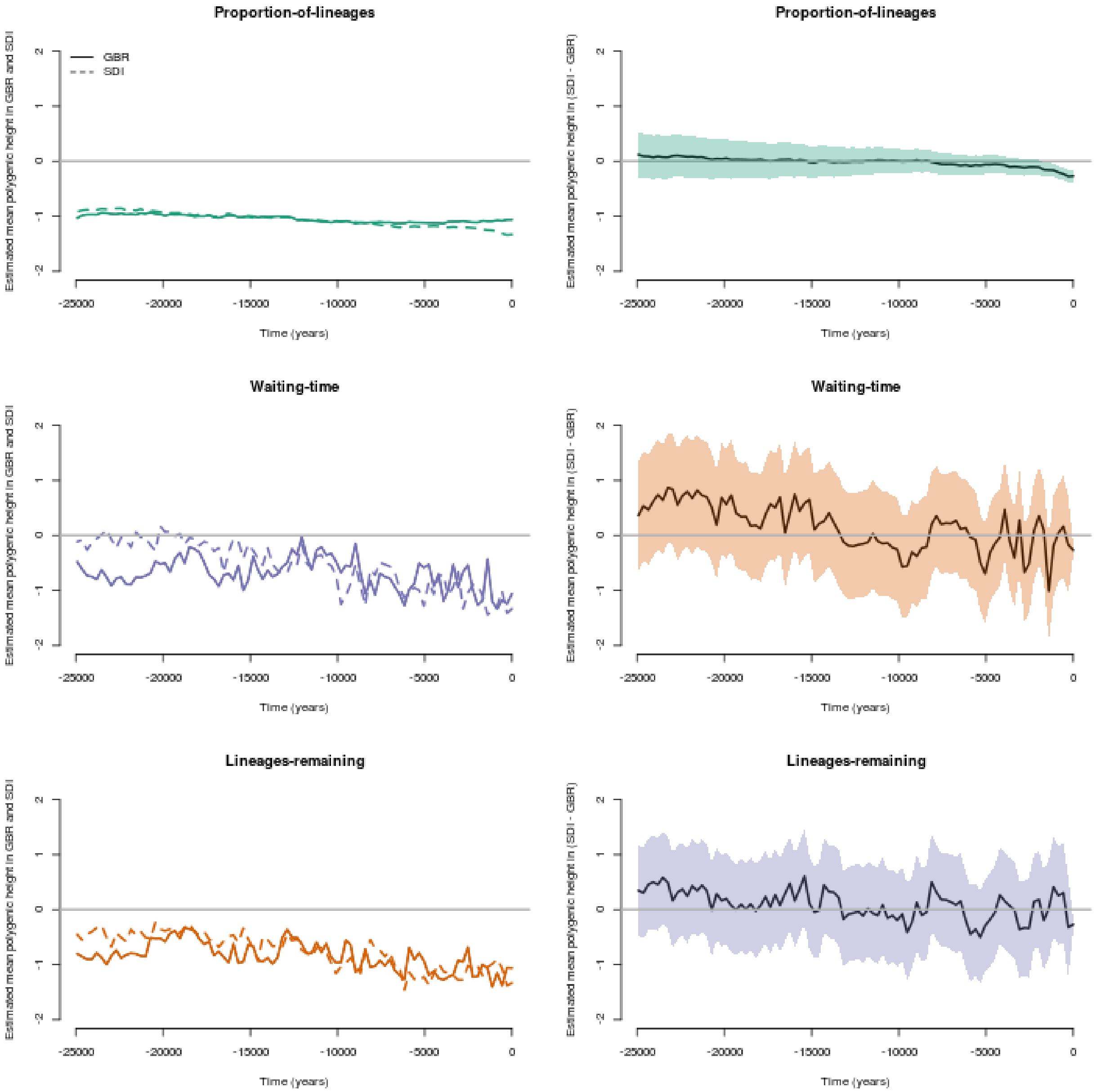


Supp Figure 3. Three estimators of mean polygenic scores for height in the populations ancestral to the GBR and SDI (Sardinians) in the past 25 ky using height loci and effect sizes ascertained from BBJ. The left column shows the mean polygenic scores in the GBR and SDI using the proportion-of-lineages estimator (the same as Figure 3), the waiting-time estimator, and the lineages-remaining estimator. The right column shows the difference between GBR and SDI in the mean polygenic scores using these estimators. Shaded areas denote the 95% confidence interval. The mean polygenic height score in Sardinia using the proportion-of-lineages estimator was significantly decreasing from that in GBR since at least 10 kya ($p = 0.0123$), while not significant between 20 kya and 10 kya ($p = 0.5307$).

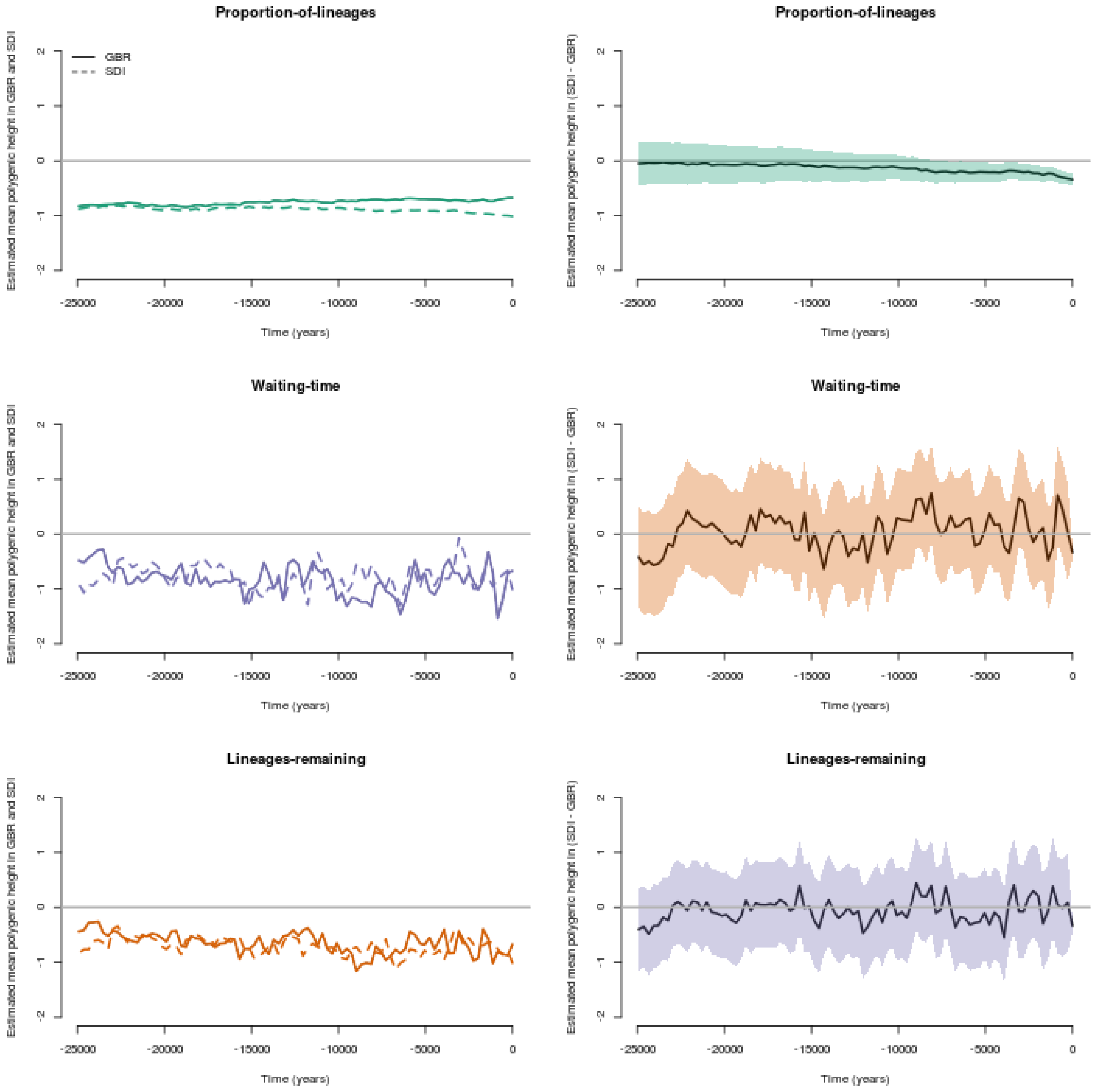


Supp Figure 4. Three estimators of mean polygenic scores for height in the populations ancestral to the GBR and SDI (Sardinians) in the past 25 ky using height loci and effect sizes ascertained from UKB. The left column shows the mean polygenic scores in the GBR and SDI using the proportion-of-lineages estimator, the waiting-time estimator, and the lineages-remaining estimator. The right column shows the difference between GBR and SDI in the mean polygenic scores using these estimators. Shaded areas denote the 95% confidence interval. The mean polygenic height score in Sardinia using the proportion-of-lineages estimator was significantly decreasing from that in GBR since at least 10 kya ($p$ = 0.0321), while not significant between 20 kya and 10 kya ($p$ = 0.5071).

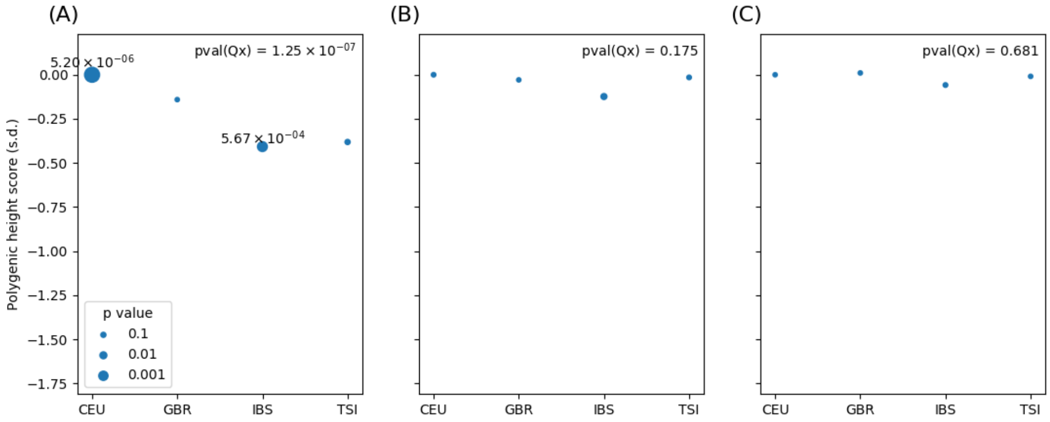


Supp Figure 5. Excess variance tests in mainland Europeans. The polygenic score was based on SNPs from (A) GIANT, (B) UKB, and (C) BBJ ascertained by picking out genome-wide significant SNPs that are more than 500 kb apart from each other. The *P* values are represented by the size of each circle, and those lower than 0.01 are shown in the plot. IBS (Iberian Population in Spain), TSI (Toscani in Italia), GBR (British in England and Scotland), and CEU (Utah Residents with Northern and Western European Ancestry) are from 1000 Genome Project.

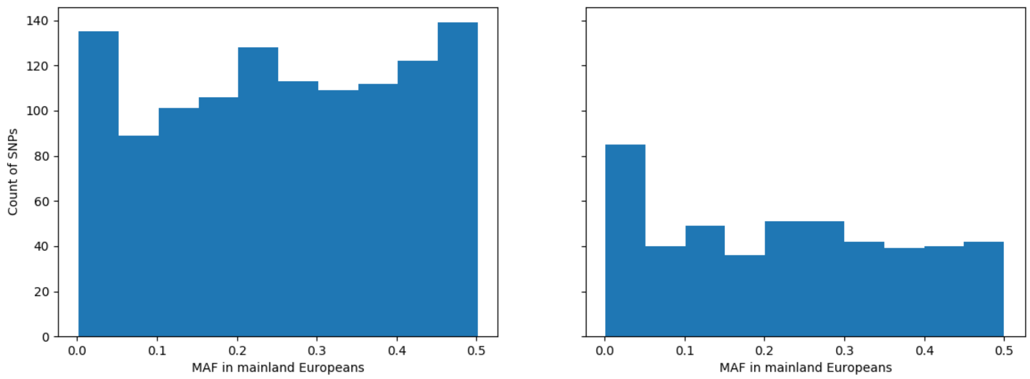


Supp Figure 6. Minor allele frequencies in mainland Europeans for SNPs ascertained from UKB (left) and BBJ (right). EUR MAF is calculated using 1000 Genomes populations CEU, GBR, IBS, and TSI.

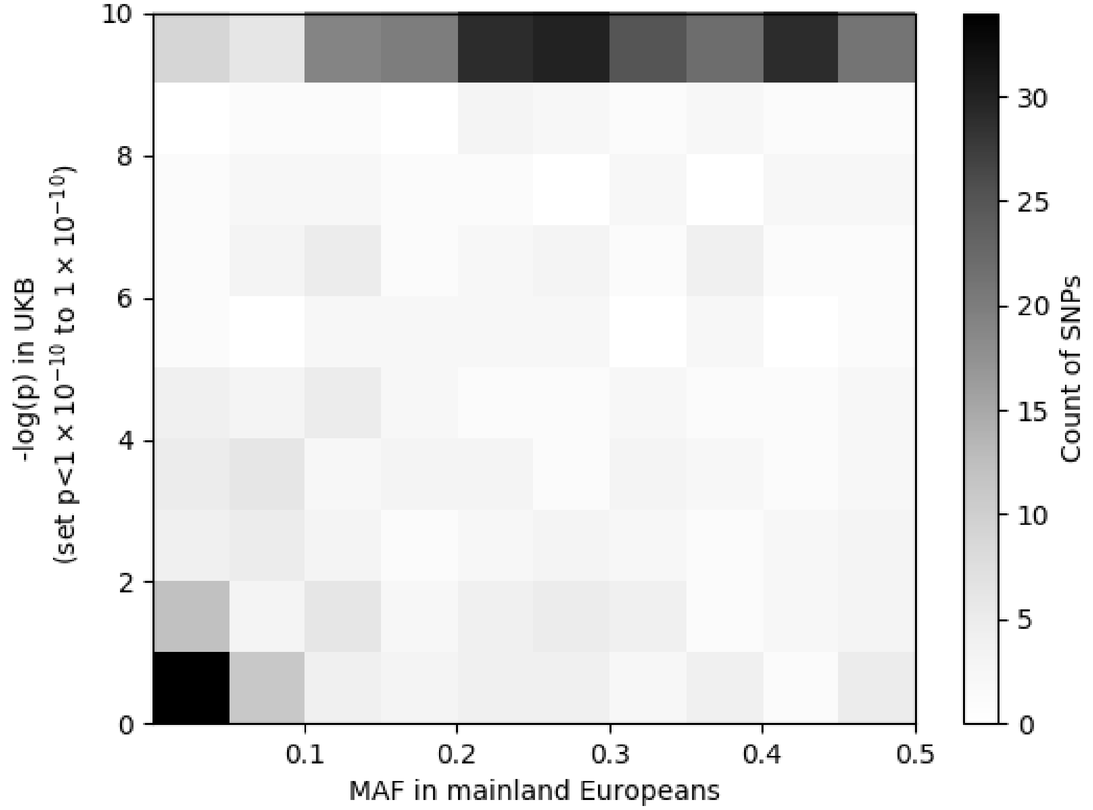


Supp Figure 7. Minor allele frequencies in European versus *p* values in UKB for SNPs ascertained from BBJ.

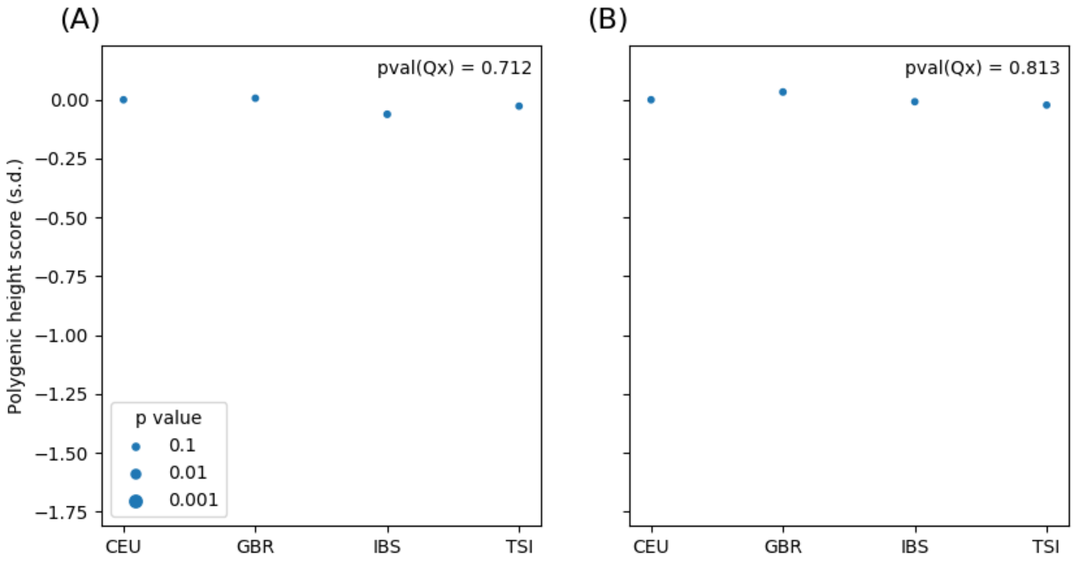


Supp Figure 8. Excess variance tests in mainland Europeans using two subsets of variants ascertained from BBJ. (A) Variants with MAF > 0.1 in mainland Europeans, and (B) variants with *p* < 5e-8 in UKB.

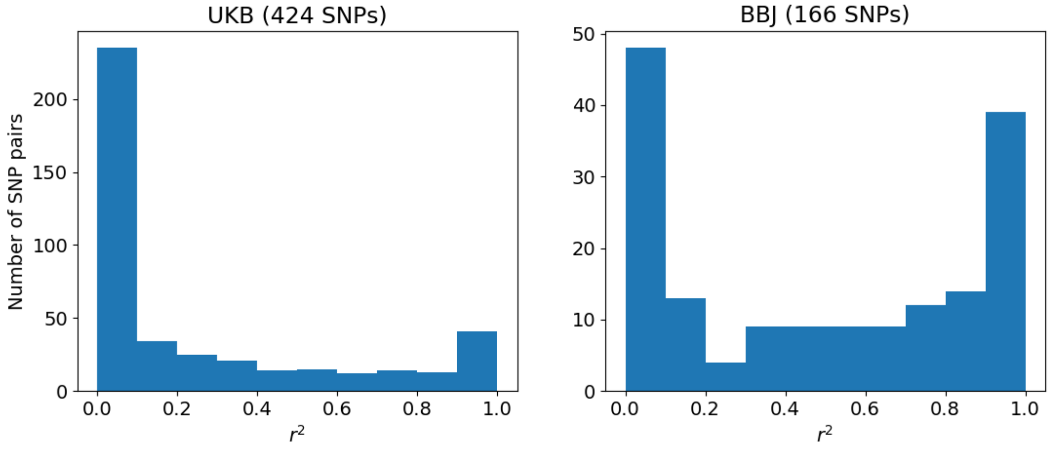


Supp Figure 9. LD in CEU between tagging, exonic, variants in gnomAD exome dataset and their nearby (less than 500 kb) most-associated, possibly non-exonic, variants in GWAS. Both the tagging and most-associated variants surpasses genome-wide significance threshold of $5x10^{-8}$.

Supp Table 1 Ascertain height-associated SNPs from BBJ but restrict to significant SNPs in UKB at the genome-wide significance threshold of 5e-8, 5e-7, and 5e-6

| GWAS p-value threshold | Mean height-increasing allele frequency difference (Northern European – Southern European) | P value from t-test |
|---|---|---|
| 5e-8 | 0.46% | 0.108 |
| 5e-7 | 0.24% | 0.332 |
| 5e-6 | 0.34% | 0.106 |

Supp Table 2 Ascertain height-associated SNPs from UKB and BBJ but restrict to SNPs which have frequencies estimated from more than 10,000 alleles in both Southern Europeans and North-Western Europeans in gnomAD exome dataset. Given the number of tests we performed and the residual stratification we expect in UKB, we caution any over-interpretation of the signals in UKB.

| GWAS panel | GWAS p-value threshold | Mean height-increasing allele difference (North-Western Europeans – Southern Europeans) | P value from t-test |
|---|---|---|---|
| UKB | 5e-8 | 0.26% | 0.063 |
| | 5e-7 | 0.25% | 0.056 |
| | 5e-6 | 0.26% | 0.029 |
| BBJ | 5e-8 | 0.16% | 0.486 |
| | 5e-7 | 0.14% | 0.496 |
| | 5e-6 | 0.08% | 0.663 |
